# Cautionary note on ribonuclease activity of recombinant PR-10 proteins

**DOI:** 10.1101/2023.02.27.529914

**Authors:** Rawit Longsaward, Nattapong Sanguankiattichai, Unchera Viboonjun, Renier A.L. van der Hoorn

## Abstract

We studied the biochemical properties of three splicing isoforms of PR-10 from rubber tree (*Hevea brasiliensis*) and found that purified recombinant *Hb*PR10 can cause RNA degradation *in vitro*, a well-known activity described for many PR-10 proteins. This ribonuclease activity was observed for all three *Hb*PR10 splicing isoforms and is abolished by boiling. However, inclusion of a negative control proteins revealed that ribonuclease activity rather originates from RNases that are copurified from *E. coli*, which are overlooked by traditionally used controls such as heat inactivation, RNase inhibitors and negative control proteins obtained with different procedures. The crucial control proteins are missing for at least nine reports on ribonuclease activity in PR-10 proteins published by different laboratories worldwide, indicating that proper controls are frequently overlooked in ribonuclease assays. The raised cautionary note applies to several PR-10 proteins with proclaimed ribonuclease activities and call for the use of different assays and mutant PR-10 proteins as control.

---

Class 10 of pathogenesis-related proteins (PR-10) is a class of PR proteins localized in the cytoplasm of the plant cell (Aglas et al., 2020). This protein family (Pfam PF00407) is originally known as a homolog of the major allergen Bet v 1 from the birch tree (Wen et al., 1997). Members of the PR-10 protein family can be induced by the infection with various pathogens including *Xanthomonas* bacteria (Bahramnejad et al., 2010), *Verticillium* fungi (Besbes et al., 2019), tobacco mosaic virus (Park et al., 2004), and sorghum mosaic virus SrMV (Peng et al., 2017). Overexpression of many PR-10 proteins increase the resistance of host plants (Zhang et al., 2021; Yang et al., 2015; Fan et al., 2015). Recently, we identified a novel PR-10 protein from rubber tree (*Hevea brasiliensis*), previously known as protein LOC110648447, which accumulates in leaves in response to root infection by white root rot fungi, *Rigidoporus microporus* (Longsaward et al., in prep). Sequence similarity and predicted 3D structure indicate that *Hb*PR10 is a member of the Bet v 1/PR-10 protein family.

The *HbPR10* gene consists of four exons that can be combined in three ways (**Figure 1A**), encoding *Hb*PR10.1, *Hb*PR10.2, and *Hb*PR10.3, which differ from each other at several residues (**Figure 1B**). To investigate *Hb*PR10s, we cloned the three splicing isoforms, synthesized by GeneArt (ThermoFisher Scientific, UK) into pET28b via Gibson assembly (Gibson et al., 2009), with a C-terminal His-tag (hexahistidine, Supplemental **Table S1**). The plasmids were confirmed by sequencing, transformed into *E. coli* Rosetta DE3 (Sigma Aldrich, USA, Supplemental **Table S2**) and selected on kanamycin and chloramphenicol. Proteins were expressed by adding 0.2 mM isopropylthio-β-galactoside (IPTG) once the culture reached OD_600_=0.6, and incubated overnight to induce the expression of recombinant proteins. Cultures were lysed in the CellLytic Express buffer (Sigma Aldrich, USA) and His-tagged proteins were purified on Ni-NTA agarose-resin (ThermoFisher, UK). The column was extensively washed and eluted with PBS buffer containing 50 and 250 mM imidazole, respectively. All three *Hb*PR10 proteins are soluble and produced and purified with high yields (**Figure 1C**).

**Figure 1.**
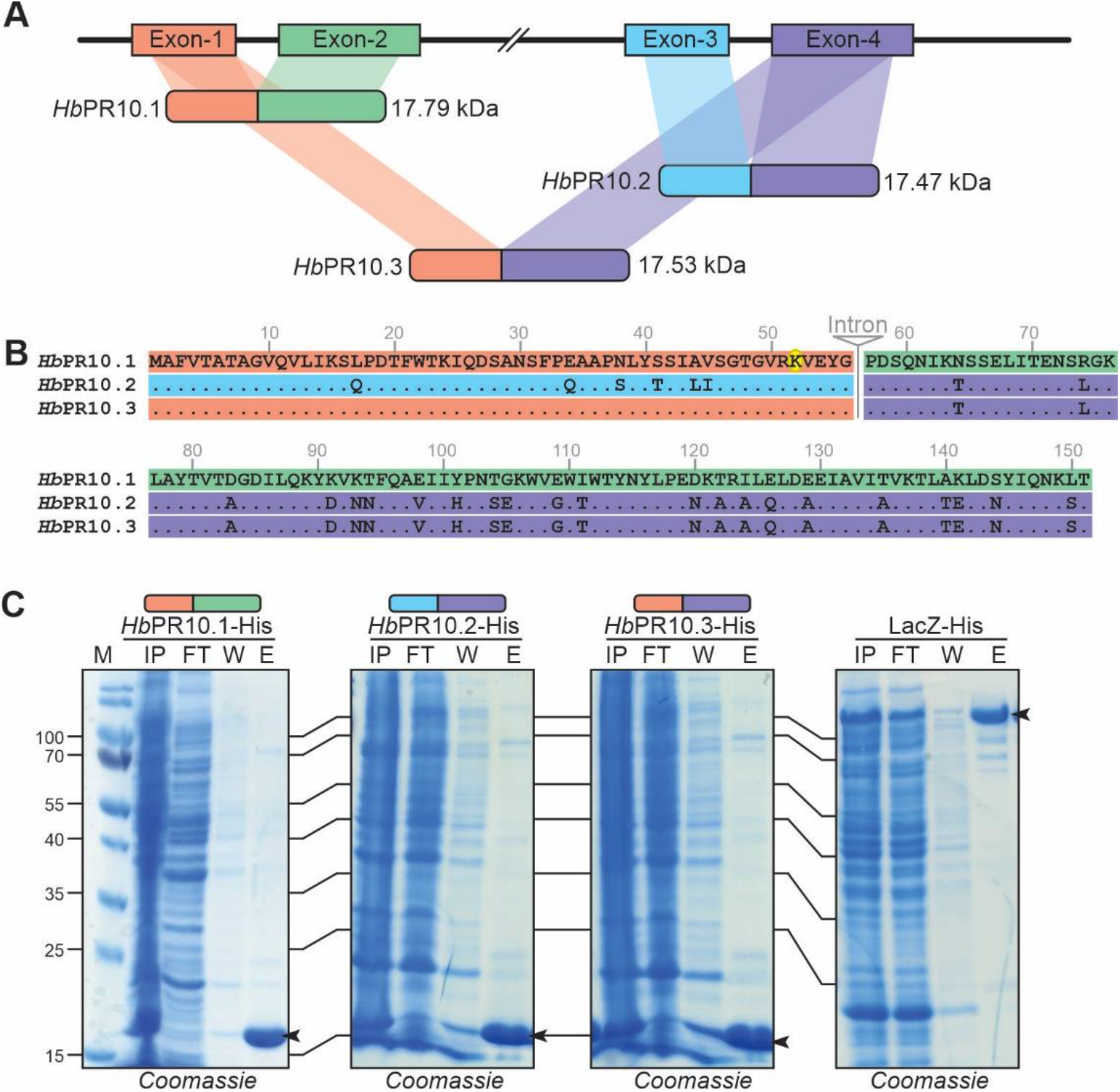
Expression and purification of three *Hb*PR-10 isoforms. **(A**) The alternative splicing among three *Hb*PR10 isoforms according to the reference sequence in the NCBI database (NW_01874736.1). The calculated molecular weights are shown. **(B**) Sequence alignment of three splicing isoforms of *Hb*PR10s. **(C)** The expression and purification of *Hb*PR10-His and LacZ-His proteins from *E. coli*. Lane IP, FT, W, and E show the protein profile from each of the Ni-NTA resin purification steps including the input, flow-through, wash, and eluate, respectively.

A frequently reported characteristic of PR-10 proteins is the ability to cleave RNA in solution, detected by separation of RNA on agarose gels (Park et al., 2004; Pungartnik et al., 2009; Huang et al., 2016). Ribonuclease activity of PR10 is thought to have a direct effect on viral RNA or on general RNA homeostasis during defence. We used this in-solution ribonuclease assay to test of *Hb*PR10s have ribonuclease activity. We incubated 1.25 μg of total RNA, extracted from *Nicotiana benthamiana* leaves via Trizol-based RNA extraction (ThermoFisher, UK) with 1 μg of purified *Hb*PR10.1-His protein for 180 min at 37 ° C, separated the RNA on 1% agarose gel and stained the gel with ethidium bromide. In the presence of *Hb*PR10.1-His, the ribosomal RNA (rRNA) was degraded and low molecular weight signals appeared (**Figure 2A**). The RNA was completely degraded by RNaseA (positive control) (**Figure 2A**). Similar ribonuclease activities were observed with purified *Hb*PR10.2-His and *Hb*PR10.3-His (**Figure 2B**).

**Figure 2.**
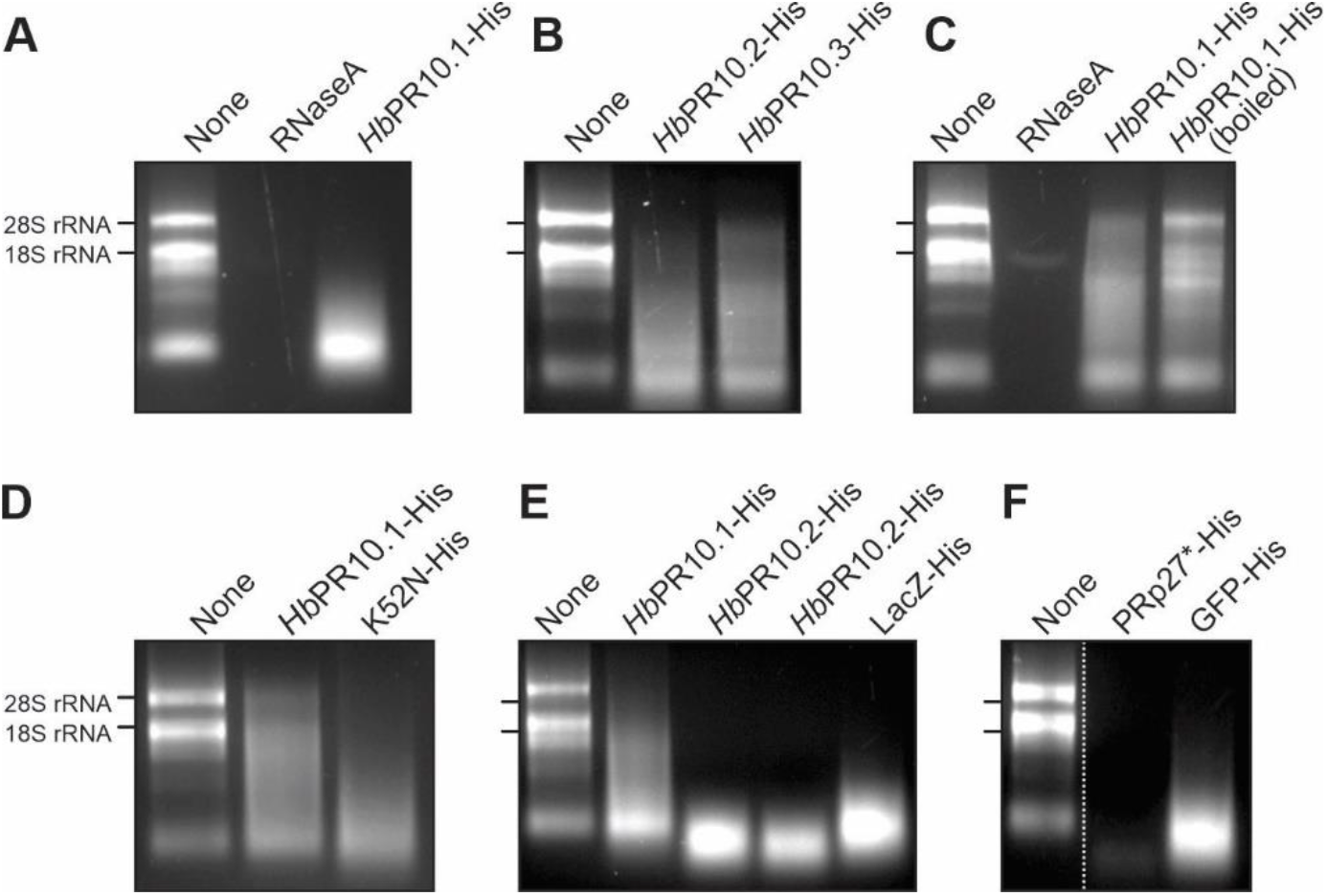
RNA degradation by purified *Hb*PR10 and other proteins. **(A**) In-solution ribonuclease activity of *Hb*PR10.1. **(B)** In-solution ribonuclease activity of *Hb*PR10.2 and *Hb*PR10.3. **(C**) In-solution ribonuclease activity is reduced upon boiling *Hb*PR10.1. **(D**) In-solution ribonuclease activity is unaltered in K52N mutant of *Hb*PR10.1. **(E**) In-solution ribonuclease activity by purified *Hb*PR10s and LacZ. **(F**) In-solution ribonuclease activity by purified PRp27 and GFP. **(A-F)** Total plant RNA (1.25 μg) was incubated with PBS buffer or purified (boiled/mutant) proteins (1 μg) at 37 ° C for 180 min, followed by 1% agarose gel electrophoresis.

Heat-inactivation is an often-used negative control in ribonuclease studies (Fan et al., 2015; Liu et al., 2006; Zubini et al., 2009; Pungartnik et al., 2009; Xie et al., 2010; Agarwal et al., 2012). Heat-treatment of *Hb*PR10-His (95 ° C for 15 min), followed by incubation with RNA resulted in a largely reduced RNA degradation compared to unheated *Hb*PR10.1-His (**Figure 2C**). We next generated the K52N substitution mutant of *Hb*PR10.1, because this mutation was found to abolish RNase activity in *Ah*PR10 (Chadha & Das, 2006). However, *Hb*PR10.1^K52N^-His is still able to degrade RNA in our in-solution assays (**Figure 2D**).

Other studies on PR-10 proteins included a protein with no reported ribonuclease activity as a negative control for the ribonuclease assay. For instance, bovine serum albumin (BSA) was included as a negative control for the in-solution ribonuclease assay of rice *Os*PR10 (Kim et al., 2008) and SPE16, a PR10 protein from jicama Mexican turnip (Wu et al., 2002). Likewise, a GST-tagged protein was included as a control for grape *Vp*PR10s (He et al., 2013; Wang et al., 2014). These negative controls, however, were not produced and purified in the same way as the PR-10 proteins. In order to confirm whether the detected ribonuclease activity was specifically from *Hb*PR10, we included a β-galactosidase with a C-terminal His-tag encoded by *LacZ*, which can cleave lactose but not RNA, as a negative control. LacZ-His was cloned into the same plasmid backbone and purified using the same methods as the three *Hb*PR10 proteins (**Figure 1C**). Alarmingly, however, purified LacZ-His also caused RNA degradation, similar to all three *Hb*PR10s (**Figure 2E**). Likewise, also GFP-His and an inactive mutant of PRp27 (PRp27^H122F^-His, Morimoto et al., 2022), expressed in *E. coli* and purified following the same procedure, was able to degrade RNA (**Figure 2F**). These experiments indicate that the ribonuclease activity may originate from some copurified proteins present in all samples.

To date, ribonuclease activity of PR-10 proteins has been investigated using three main assays (**Figure 3**, Supplemental **Table S3**): i) in-solution ribonuclease assay followed by agarose gel electrophoresis; ii) in-solution ribonuclease assay coupled to spectrophotometry at the OD_260_ to calculate the unit of enzyme activity; and iii) ingel ribonuclease assay using polyacrylamide gel electrophoresis co-polymerized with RNA, which was first described by Yen and Green (1991). Based on the results we obtained here, *Hb*PR10 proteins are unproven to have ribonuclease activity at the conditions tested because we detected ribonuclease activity also for the negative control proteins. We found that Nickel affinity column for purification cannot eliminate copurified contaminant ribonucleases and hence cause RNA degradation, even by negative control proteins. Even though the in-solution assay is the easy and frequently used (**Figure 3**, Supplemental **Table S3**), copurified ribonucleases contaminating the eluates are easily overlooked (**Figure 2**). Inactivation by boiling or by using ribonuclease inhibitors is insufficient as a negative control, and also the use of proteins that were not produced and purified following the same procedure are unsuitable controls. Consequently, previous conclusions that *Ss*PR10 (Liu et al., 2006), Pru p s (Zubini et al., 2009), *Tc*PR-10 (Pungartnik et al., 2009), *Zm*PR10.1 (Xie et al., 2010), *ann*PR10 and *bac*PR10 (Soh et al., 2012), *Jc*PR-10a (Agarwal et al., 2013), Gly m 41 (Fan et al., 2015), *Pn*PR-like (Li et al., 2021), *Ma*PR10s (Rajendram et al., 2022) have ribonuclease activities should be reconsidered given the absence of proper controls.

**Figure 3.**
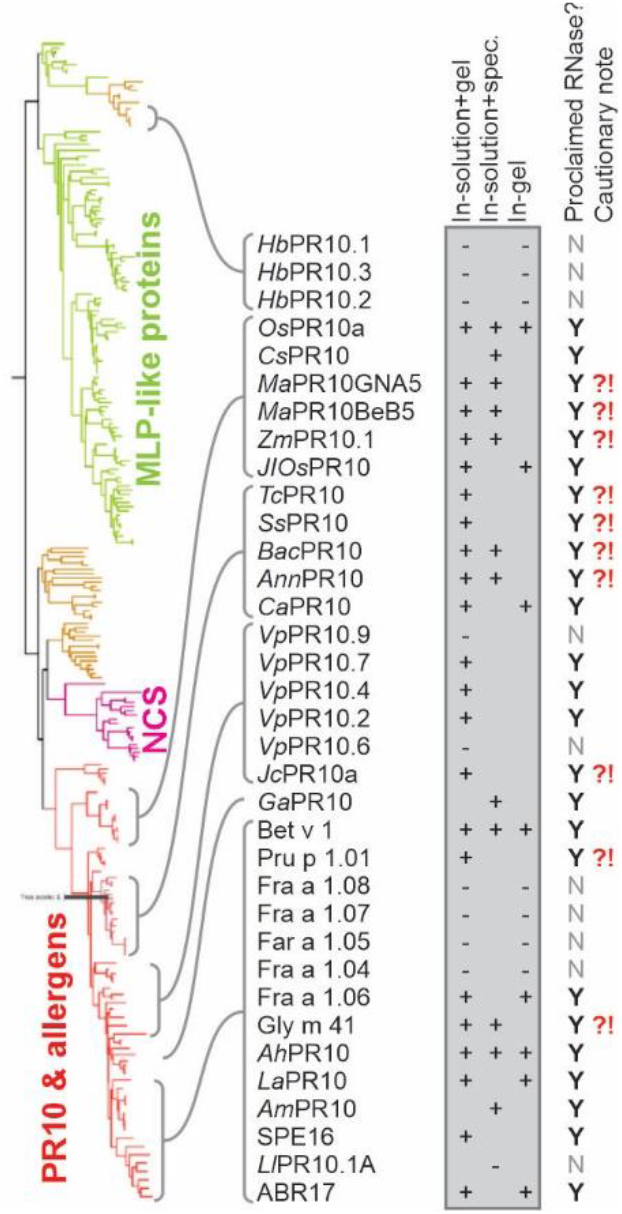
Caution on proclaimed ribonuclease activities of PR10 homologs. PR10 homologs studied in ribonuclease assays are ranked phylogenetically. Ribonuclease assays are: 1) in-solution ribonuclease assay, followed by agarose gel electrophoresis; 2) in-solution ribonuclease assay, followed by spectrometric analysis; 3) in-gel ribonuclease assay. Ribonuclease activity was observed (+) or not (−), resulting in proclaimed ribonuclease activities (Y). Proteins with proclaimed ribonuclease activity but insufficient controls are noted with red question/caution mark (**?!**). The Maximum Likelihood tree was generated with the PhyML server.

The absence of ribonuclease activity for *Hb*PR10 is consistent with other reported PR-10 proteins that lack ribonuclease activity such as *Vp*PR10.6 and *Vp*PR10.9 (Wang et al., 2014), *Ll*PR-10.1 A (Biesiadka et al., 2002), and Fra a 1 proteins (Besbes et al., 2019). The construction of an evolutionary tree with reported PR-10 protein homologs (**Figure 3**) showed that potentially false positive ribonuclease activities are common across the PR-10 subfamily. *Hb*PR10 proteins are members of the major-latex protein (MLP) subgroup of the PR-10 family. No MLP proteins are reported with ribonuclease activity (Fujita and Inui, 2021), which is consistent with our results (**Figure 3**). We hope this cautionary note will encourage the community to develop appropriate assays and include adequate negative controls to detect ribonuclease activity. Given the sensitivity for RNA degradation by contaminant RNAses, mutant PR-10 proteins that lack ribonuclease activity but are produced and purified in the same way are essential negative controls for future assays.

## Acknowledgements

This research was financially supported by the Science Achievement Scholarship of Thailand (SAST. RL); BBSRC 19RM3 project ‘Galactosyrin’ BB/T015128/1 (NS) and the European Research Council ERC-AdG-2020 project 101019324 ‘ExtraImmune’ (RvdH).

## Competing interests

The authors declare that they have no competing interests.

## Availability of data and materials

All data generated or analysed during this study are included in this published article and its supplementary information file.

## Notes

### Competing Interest Statement

The authors have declared no competing interest.

