## Supplemental Table for "Cautionary note on ribonuclease activity of recombinant PR-10 proteins"

### Supplemental Tables S1-S2-S3

### Cautionary note on ribonuclease activity of recombinant PR-10 proteins

Rawit Longsaward, Nattapong Sanguankiattichai, Unchera Viboonjun, Renier A.L. van der Hoorn

**Table S1** Oligonucleotides used in this study

| Fragment | Primer name | Sequence (5' to 3') |
|---|---|---|
| *Hb*PR10.1-His | PR10_pET28b_F | CCCTCTAGAAATAATTTTGTTTAACTTTAAGAAGGAGATATACAATGGCTTTCGTGACTGCTACTGCT |
| | PR10_pET28b_R1 | GTTAGCAGCCGGATCCAAGCCTAGTGATGGTGATGGTGATGGGTCAGCTTGTTCTGGATGTAG |
| *Hb*PR10.2-His | PR10_pET28b_F | CCCTCTAGAAATAATTTTGTTTAACTTTAAGAAGGAGATATACAATGGCTTTCGTGACTGCTACTGCT |
| | PR10_pET28b_R2 | GTTAGCAGCCGGATCCAAGCCTAGTGATGGTGATGGTGATGGGTGCTCTTGTTCTGGATGTAG |
| *Hb*PR10.3-His | PR10_pET28b_F | CCCTCTAGAAATAATTTTGTTTAACTTTAAGAAGGAGATATACAATGGCTTTCGTGACTGCTACTGCT |
| | PR10_pET28b_R2 | GTTAGCAGCCGGATCCAAGCCTAGTGATGGTGATGGTGATGGGTGCTCTTGTTCTGGATGTAG |
| His-GFP-Strep | oNS368 | CCCTCTAGAAATAATTTTGTTTAACTTTAAGAAGGAGATATACAATGGCGCATCACCATCACCATC |
| | oNS369 | GTTAGCAGCCGGATCCAAGCTCACTTTTCGAACTGCGGGTGGCTCCAGCTACCTTTGTACAGTTCATCCATACCATGCG |
| LacZ-His | oNS339 | CCCTCTAGAAATAATTTTGTTTAACTTTAAGAAGGAGATATACAATGACCATGATTACGGATTCAC |
| | oNS340 | GTTAGCAGCCGGATCCAAGCTTAGTGATGGTGATGGTGATGTTTTTGACACCAGACCAACT |

**Table S2** Plasmids used in this study

| Plasmid | Description | Reference |
|---|---|---|
| pRL010 | *Hb*PR10.1-His in pET28b vector | *This work* |
| pRL011 | *Hb*PR10.2-His in pET28b vector | *This work* |
| pRL012 | *Hb*PR10.3-His in pET28b vector | *This work* |
| pNS141 | LacZ-His in pET28b vector | *This work* |
| pNS153 | His-GFP-Strep in pET28b vector | *This work* |
| pKM008 | His-PRp27(H122F) | Morimoto et al., 2022 |

**Table S3** Ribonuclease assays on PR-10 protein used in previous studies.

| | | | | | | Downstream process | | | RNase assay | | | Negative controls | | | | | | | |
|---|---|---|---|---|---|---|---|---|---|---|---|---|---|---|---|---|---|---|---|
| Types | Protein | Accession no. | Original species | Fusion tag | Purification method | Dialyzed | Fractioned | Remove tag | In-solution and agarose gel electrophoresis | In-solution and spectrophotometry | In-gel ribonuclease | Boiled protein | Mutant protein | RNase inhibitor | Buffer (no protein) | (+) DTT | Empty vector | Other protein | Reference |
| | *Ca*PR10 | AAF63519.1 | *Capsicum annuum* | - | Crude protein extraction | - | - | - | + | - | + | - | - | + | + | + | + | - | Park et al. 2004 |
| Native | SPE16 | ARR11455.1 | *Pachyrrhizus erosus* | - | DEAE-Sepharose column | - | + | - | + | - | - | - | - | - | + | - | - | + | Wu et al. 2002 |
| Native | *Os*PR10a | XP_015620382.1 | *Oryza sativa* | - | Continuous-elution electrophoresis | - | + | - | + | + | + | - | - | - | + | - | - | - | Huang et al. 2016 |
| Native | Bet v 1 | 1FM4_A | *Betula alba* | - | Fast protein liquid chromatography (FPLC) | + | + | - | + | + | + | - | - | + | + | - | - | - | Bufe et al. 1996 |
| Native | *Am*PR10 | ASA69247.1 | *Astragalus mongholicus* | - | Phosphate buffer | + | + | - | - | + | - | - | - | - | - | - | - | - | Yan et al. 2008 |
| Native | Bet v 1 | 1FM4_A | *Betula alba* | - | Reverse phase HPLC | - | - | - | + | - | + | - | - | - | + | - | - | - | Swoboda et al. 1996 |
| Native | *As*PRs | - | *Angelica sinensis* | - | Sephadex G50; Ion exchange | | | | - | + | - | - | - | - | - | - | - | - | Pan et al. 2018 |
| Recombinant | Fra a 1s | AHZ10955.1, AHZ10956.1, AHZ10957.1, AHZ10958.1, AHZ10959.1 | *Fragaria x ananassa* | C-term hexahistidine | Affinity purified | - | - | - | + | - | + | + | - | - | + | - | + | - | Besbes et al. 2019 |

| | | | | | | | | | | | | | | | | | | | |
|---|---|---|---|---|---|---|---|---|---|---|---|---|---|---|---|---|---|---|---|
| Recombinant | *Cs*PR10 | ADL09408.1 | *Crocus sativus* | GST tag | GS-4B resin, followed by MALDI analysis | - | - | - | - | + | - | - | - | - | - | + | - | - | Gomez-Gomez et al. 2011 |
| Recombinant | *Ann*PR10 | ABC74798.1 | *Capsicum annuum* | GST tag | GST tag purification | - | - | - | + | + | - | - | - | - | + | - | - | - | Soh et al. 2012 |
| Recombinant | *Bac*PR10 | ABC74797.1 | *Capsicum baccatum* | GST tag | GST tag purification | - | - | - | + | + | - | - | - | - | + | - | - | - | Soh et al. 2012 |
| Recombinant | *Vp*PR10.2 | ABD78556.1 | *Vitis pseudoreticulata* | GST tag | GST tag purification | - | - | - | + | - | - | + | - | + | - | - | - | + | He et al. 2013 |
| Recombinant | *Vp*PR10s | - | *Vitis pseudoreticulata* | GST tag | GST tag purification | - | - | - | + | - | - | + | - | - | + | - | - | + | Wang et al. 2014 |
| Recombinant | Gly m 41 | ADX43926.1 | *Glycine max* | Hexahistidine | His-bind resin column | - | - | - | + | + | - | + | - | - | + | - | - | - | Fan et al. 2015 |
| Recombinant | *Ss*PR10 | AAU00066.1 | *Solanum surattense* | Polyhistidine | His-bond Ni Affinity resin column | + | - | - | + | - | - | + | - | - | + | - | + | - | Liu et al. 2006 |
| Recombinant | SPE16 | ARR11455.1 | *Pachyrrhizus erosus* | n/a | Ni-chelating Sepharose Fast Flow Gel column | - | - | - | + | - | - | - | + | - | + | - | - | - | Wu e al. 2003 |
| Recombinant | *Zm*PR10.1 | ADA68331.1 | *Zea mays* | C-term hexahistidine | Ni-IDA affinity column | - | - | - | + | + | - | + | - | + | - | - | + | - | Xie et al. 2010 |
| Recombinant | ABR17 (PR10.4) | Q06931.1 | *Pisum saivum* | N-term hexahistidine | Ni-NTA agarose column | + | - | + | + | - | + | + | - | - | - | - | - | - | Srivastava et al. 2006a, 2007 |
| Recombinant | ABR17 (PR10.4) | Q06931.1 | *Pisum saivum* | N-term hexahistidine | Ni-NTA agarose column | + | - | - | + | - | - | + | + | - | - | - | - | - | Krishnaswamy et al. 2011 |
| Recombinant | *Ca*PR10 | AAF63519.1 | *Capsicum annuum* | N-term hexahistidine | Ni-NTA agarose column | - | - | - | + | - | - | + | - | - | + | - | + | - | Park et al. 2004 |
| Recombinant | *Ga*PR-10 | AAL09033.1 | *Gossypium arboreum* | N-term hexahistidine | Ni-NTA agarose column | - | - | + | - | + | - | - | - | - | - | - | - | - | Zhou et al. 2002 |
| Recombinant | *Jc*PR-10a | AEV54115.1 | *Jatropa curcas* | Hexahistidine | Ni-NTA agarose column | - | - | - | + | - | - | + | - | - | + | - | - | - | Agarwal et al. 2012 |
| Recombinant | *JIOs*PR10 | AAL74406.1 | *Oryza sativa* | C-term hexahistidine | Ni-NTA agarose column | - | - | - | + | - | + | - | + | + | + | + | - | + | Kim et al. 2008 |
| recombinant | *La*PR-10 | CAA03926.1 | *Lupinus albus* | C-term hexahistidine | Ni-NTA agarose column | - | - | - | + | - | + | - | - | + | - | - | - | - | Bantignies et al. 2000 |
| Recombinant | Pea PR10.1 | - | *Pisum saivum* | N-term hexahistidine | Ni-NTA agarose column | + | - | + | + | - | + | + | - | - | + | - | - | - | Srivastava et al. 2006b |

| | | | | | | | | | | | | | | | | | | | |
|---|---|---|---|---|---|---|---|---|---|---|---|---|---|---|---|---|---|---|---|
| Recombinant | *Pn*PR-like | QOJ53932.1 | *Panax notoginseng* | Polyhistidine | Ni-NTA agarose column | - | - | - | + | - | - | - | - | + | + | - | - | - | Li et al. 2021 |
| Recombinant | Pru p 1s | ACE80940.1 | *Prunus persica* | N-term hexahistidine | Ni-NTA agarose column | + | - | - | + | - | - | + | - | - | + | - | + | - | Zubini et al. 2009 |
| Recombinant | *Ma*PR10s | UED15064.1, UED15065.1 | *Musa acuminata* | N-term hexahistidine | Nickle His Gravitrap affinity column | - | - | - | + | + | - | - | - | - | + | - | - | - | Rajendram et al. 2022 |
| Recombinant | Bet v 1 | 1FM4_A | *Betula alba* | - | PBE-94 exchange column | - | + | - | - | - | + | - | - | - | + | - | - | - | Swoboda et al. 1996 |
| Recombinant | *Ah*PR10 | AAU81922.1 | *Arachis hypogaea* | C-term hexahistidine | Talon resin metal affinity column | - | + | - | + | + | + | - | + | - | + | - | - | - | Chadha and Das 2006 |
| Recombinant | *Tc*PR-10 | - | *Theobroma cacao* | N-term hexahistidine | Talon resin metal affinity column | + | - | - | + | - | - | + | - | - | + | - | - | - | Pungartnik et al. 2009 |

Note: "+" indicates the method used in the study and "-" indicates the method was not used in the study.
